## Supplementary Figures for "Cellular mechanisms of heterogeneity in *NF2*-mutant schwannoma"

**Supplementary Figure 1.** Enhanced multipolarity and cytoskeletal responses in *Nf2*-deficient Schwann cells; related to Figure 1.

**Supplementary Figure 2**. Adhesion and signaling in *Nf2*$^{-/-}$ SCs in the presence of Nrg1; related to Figure 2.

**Supplementary Figure 3.** Adhesion and signaling in *Nf2*$^{-/-}$ SCs in the presence of Nrg1; related to Figure 2.

**Supplementary Figure 4.** *Nf2*$^{-/-}$ SCs upregulate basal polarity components upon Nrg1-deprivation; related to Figure 3.

**Supplementary Figure 5**. *Nf2*$^{-/-}$ SCs enact a distinct feedforward autocrine Nrg1 signaling program upon nutrient deprivation; related to Figure 4.

**Supplementary Figure 6.** Heterogeneous mTOR activation and drug sensitivity; related to Figure 5.

**Supplementary Figure 7.** Self-generated heterogeneity; related to Figure 6.

**Supplementary Figure 8.** Quantitative analysis of self-generated heterogeneity in human schwannoma tissue; related to Figure 7.

**Supplementary Figure 9.** Quantitative analysis of heterogeneity in schwannomas arising in *Postn-Cre;Nf2*$^{lox/lox}$ DRGs; related to Figure 8.

### Supplementary Files

Supplementary Movie 1.

Supplementary Movie 2.

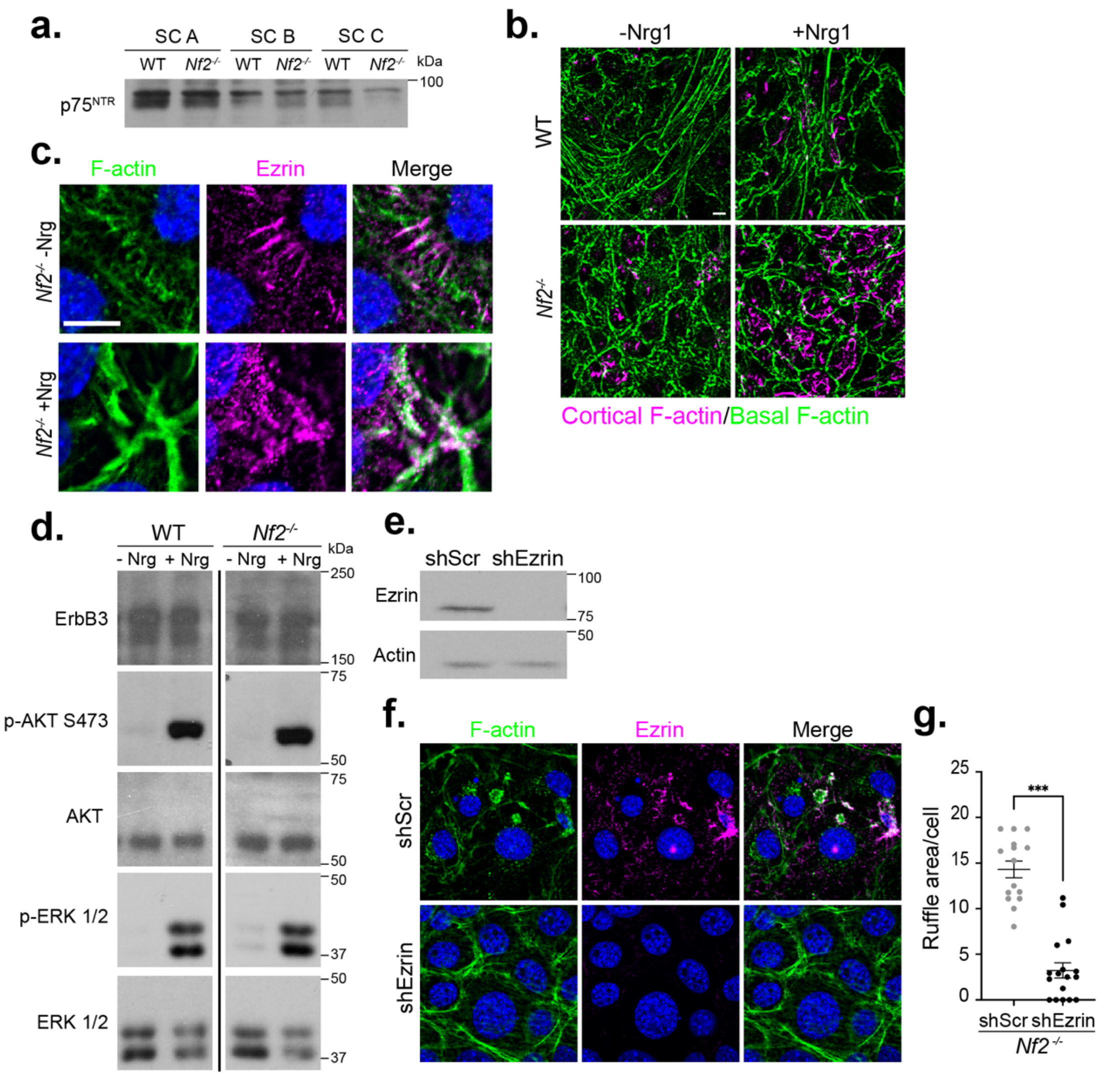


**Supplementary Figure 1. A**. Representative immunoblot showing protein levels of the SC marker p75 in *Nf2^flox/flox^* (WT) and *Nf2*$^{-/-}$ SCs. **B.** Confocal images depicting enhanced Nrg1-induced cortical ruffling in late confluent *Nf2*$^{-/-}$ but not control SCs. Images were generated by overlaying MIP of cortical actin (magenta) and basal actin (green) from 3D z stack images. **C.** Confocal images highlighting ezrin (magenta) and F-actin (green) distribution to microvilli (-Nrg1) and cortical ruffles (+Nrg1). **D.** Representative immunoblot showing protein levels of ErbB3 and activation of pAKT S473 and pERK1/2 in response to stimulation of WT and *Nf2*$^{-/-}$ SCs with Nrg. Total levels of AKT and ERK1/2 served as loading controls. The line between lanes 2 and 3 indicates the removal of unrelated lanes from the image of the blot. **D.**

Representative immunoblot showing protein levels of ezrin in *Nf2*$^{-/-}$ SCs infected with shSCR- or shEzrin-expressing lentiviruses. Actin served as a loading control. **E.** Confocal images showing ezrin (magenta) and F-actin (green) localization in *Nf2*$^{-/-}$ SCs infected with shSCR or shEzr-expressing lentiviruses. **F.** Quantitation of cortical ruffling area in *Nf2*$^{-/-}$ SCs infected with shSCR or shEzr-expressing lentiviruses. Data points are shown for n = 15 cells per condition. Data are presented as mean +/- SEM. N = 2 independent experiments. ***p<0.001, unpaired two-tailed Student's t-test. Scale bars = 10 μm.

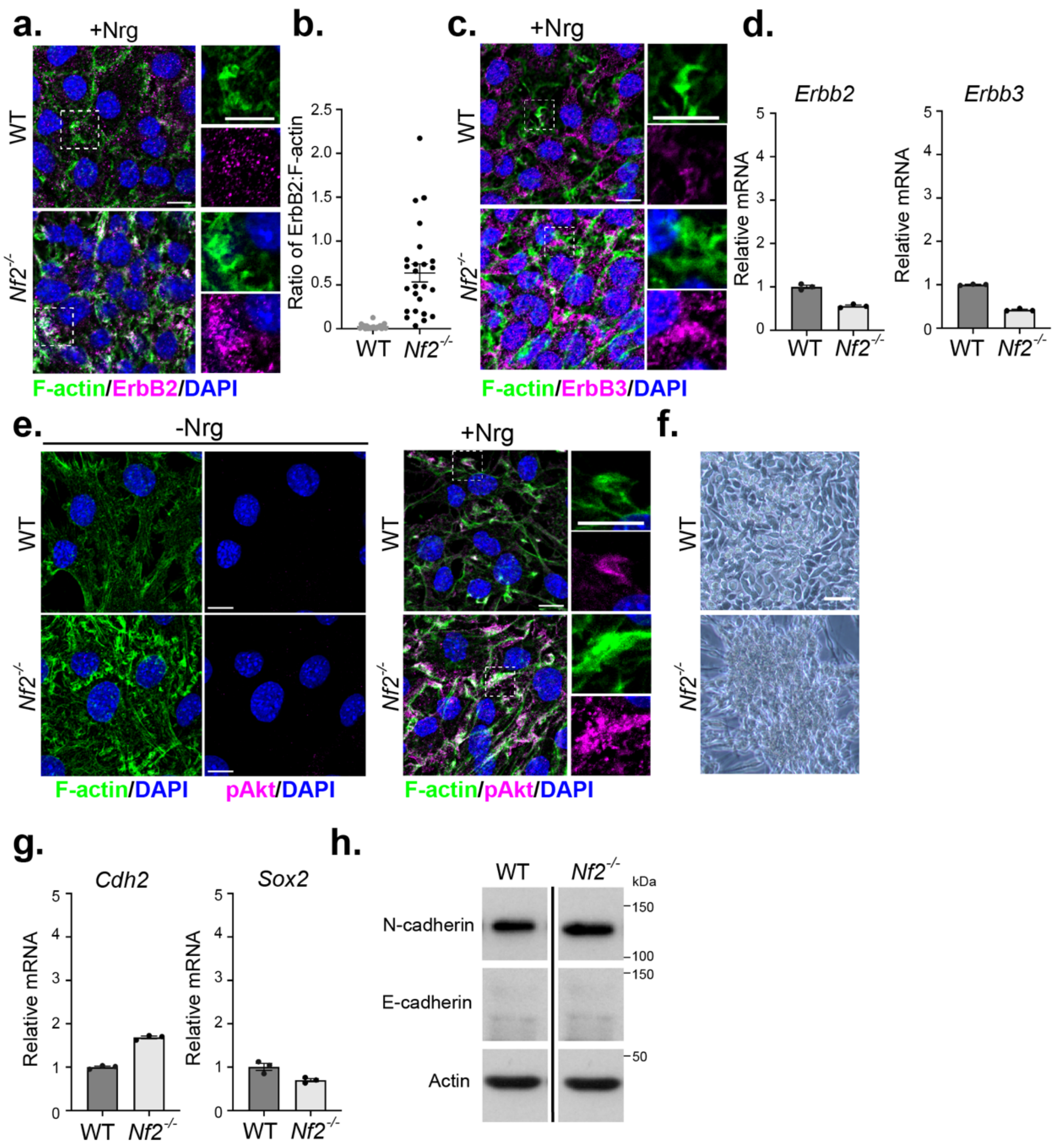


**Supplementary Figure 2**. **A.** Confocal images showing recruitment of ErbB2 (magenta) to Nrg1-induced F-actin (green)-enriched cortical ruffles in WT and *Nf2*$^{-/-}$ SCs. **B.** Quantitation of the ratio of ErbB2 relative to F-actin in Nrg1-induced ruffles in WT and *Nf2*$^{-/-}$ SCs. Data points are shown for n = 20 (WT) or n = 25 (*Nf2*$^{-/-}$) cells. N = 3 independent experiments. **C.** Confocal images of Nrg1-stimulated *Nf2*$^{-/-}$ SCs labelled for ErbB3 (magenta) and F-actin (green) to show recruitment of ErbB3 to cortical ruffles. **D**. Measurement of ErbB2 and ErbB3 mRNA levels in WT and *Nf2*$^{-/-}$ SCs by qPCR. Bars represent mean +/- SEM. **E.** Confocal images showing pAkt

(S473) (magenta) and F-actin (green) distribution in *Nf2*$^{-/-}$ SCs with or without Nrg1 stimulation. **F.** Phase contrast images of late confluent WT and *Nf2*$^{-/-}$ SCs. **G.** mRNA expression of *Sox2* and N-cadherin (*Cdh2*) in WT and *Nf2*$^{-/-}$ SCs. Data is presented as mean +/- SEM relative to WT mRNA levels. **H.** Immunoblot showing protein levels of N-cadherin and E-cadherin in WT and *Nf2*$^{-/-}$ SCs. Actin served as a control and was run with the same samples in parallel on a separate blot. The line between lanes indicates the removal of unrelated lanes from the image of the blot. Scale bars = 10 μm.

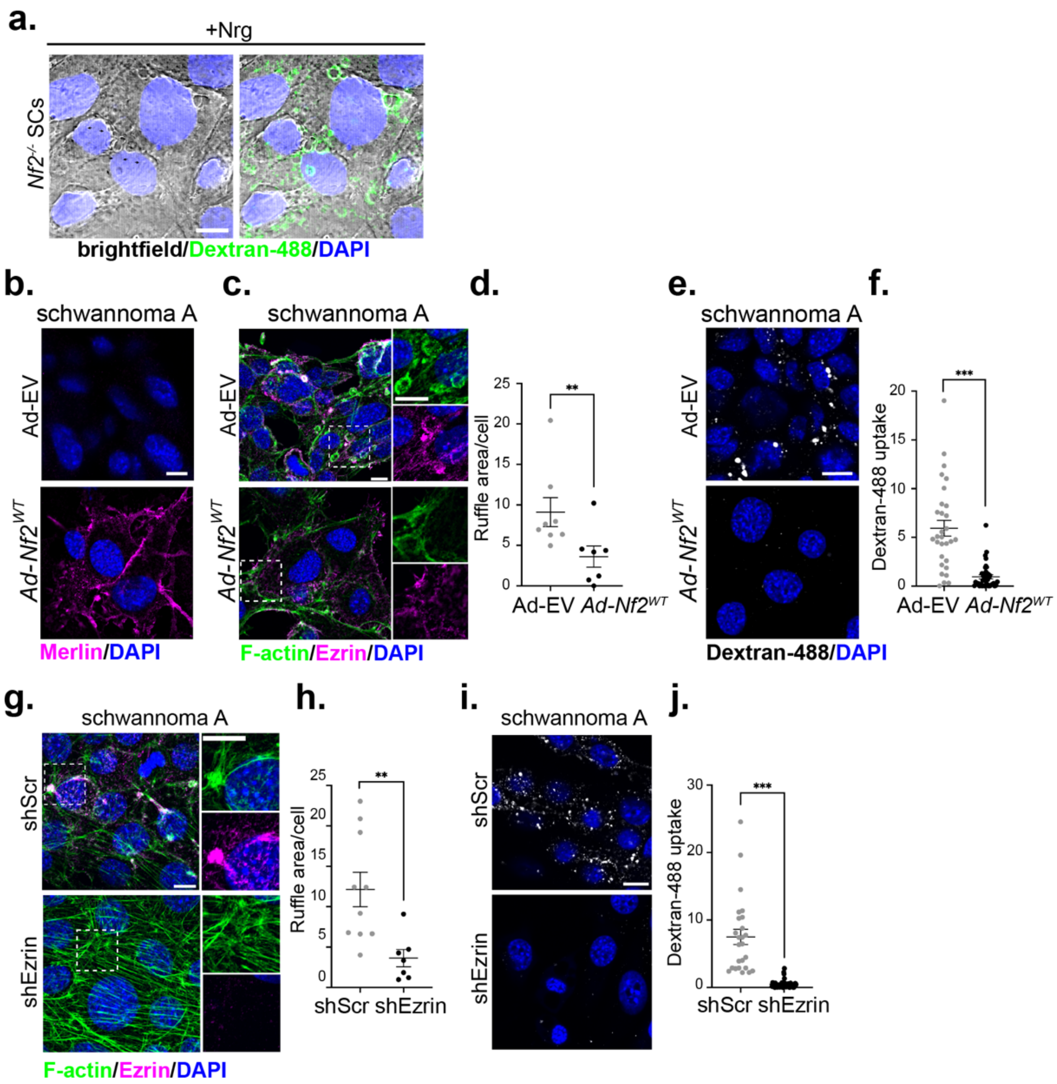


**Supplementary Figure 3. A.** Brightfield images with and without overlay of dextran-488 labelling showing uptake in large macropinosome-like structures in Nrg1-stimulated *Nf2*$^{-/-}$ SCs. **B.** Confocal images showing merlin (magenta) localization in *Nf2*$^{-/-}$ schwannoma cells expressing EV or *Nf2*$^{WT}$ via adenoviral infection. **C.** Confocal images of F-actin (green) and ezrin (magenta) localization in *Nf2*$^{-/-}$ schwannoma cells expressing EV or *Nf2*$^{WT}$ adenovirus. **D.** Quantitation of cortical ruffling in cells shown in (B). Data points are shown for n = 8 (Ad-EV) or n = 7 (*Ad-Nf2*$^{WT}$) cells. N = 2 independent experiments. **E.** Dextran-488 uptake in *Nf2*$^{-/-}$ schwannoma cells expressing EV or *Nf2*$^{WT}$ adenovirus. **F.** Quantitation of data from (E). Data points are shown as n = 30 (Ad-EV) or n = 39 (*Ad-Nf2*$^{WT}$) cells. N = 2 independent experiments.

**G.** Confocal images of F-actin (green) and ezrin (magenta) localization in *Nf2*$^{-/-}$ schwannoma cells infected with shSCR- or shEzrin-expressing lentivirus. **H.** Quantitation of cortical ruffling of cells in (G). Data points are shown for n = 10 (shScr) or n = 7 (shEzrin) cells. N = 2 independent experiments. **I.** Dextran-488 uptake in *Nf2*$^{-/-}$ schwannoma cells expressing shSCR- or shEzrin lentivirus. **J.** Quantitation of data from (I). Data points are shown for n = 25 (shScr) or n = 42 (shEzrin) cells. N = 2 independent experiments. For all graphs, lines represent mean +/- SEM. **p<0.01, ***p<0.001, unpaired two-tailed Student's t-test. Scale bars = 10 μm.

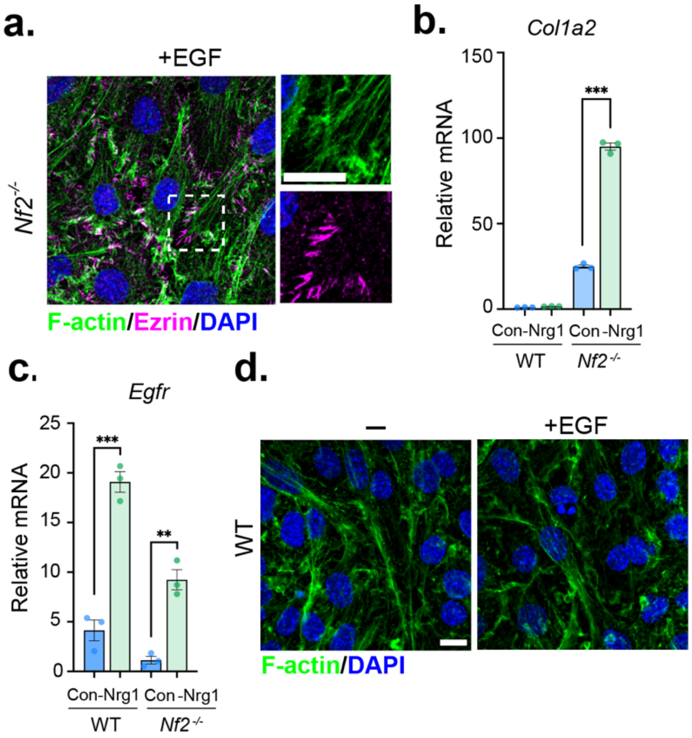


**Supplementary Figure 4. A.** Confocal images depicting the distribution of F-actin (green) and ezrin (magenta) in *Nf2*-/- SCs starved of Nrg1 overnight and stimulated with 10 ng/ml EGF for 10 min. **B.** Levels of *Col1a2* mRNA in Con. or Nrg-deprived WT or *Nf2*-/- SCs. Data is presented as mean +/- SEM relative to control, WT mRNA levels for n = 3 replicates. **C.** Levels of *Egfr* mRNA in Con. or Nrg-deprived WT or *Nf2*-/- SCs. Data is presented as mean +/- SEM relative to control, WT mRNA levels for n = 3 replicates. **D.** Confocal images of F-actin (green) localization in WT SCs starved of Nrg1 overnight and stimulated with EGF. **p<0.01, ***p<0.001, unpaired two-tailed Student's t-test. Scale bars = 10 μm.

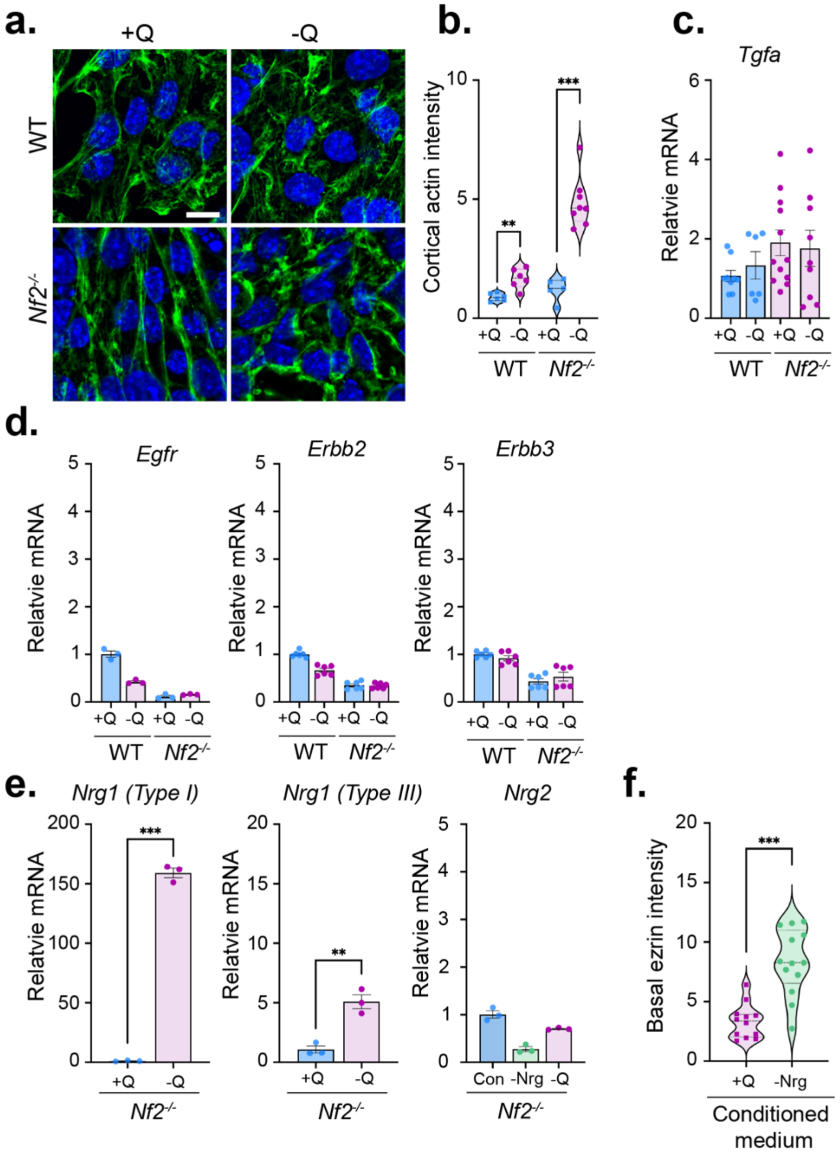


**Supplementary Figure 5**. **A.** Confocal images depicting F-actin (green) localization in WT and *Nf2*$^{-/-}$ SCs grown in complete medium (+Q) or glutamine-deprived medium (-Q) for 24 h. Scale bar = 10 μm. **B.** Quantitation of cortical ruffling in cells shown in (A). Data points are shown for n = 6 (WT, +Q), n = 6 (WT, -Q), n = 5 (*Nf2*$^{-/-}$, +Q), or n = 42 (*Nf2*$^{-/-}$, -Q) cells. N = 3 independent experiments. **C.** *Tgfa* mRNA levels in WT and *Nf2*$^{-/-}$ SCs grown in complete (+Q) or glutamine-deprived (-Q) medium for 24 h. Data is presented as mean +/- SEM relative to WT, +Q mRNA levels for n = 3 replicates. N = 3 independent experiments. **D**. *Egfr*, *ErbB2* and *ErbB3* mRNA expression in WT and *Nf2*$^{-/-}$ SCs grown in complete (+Q) or glutamine-deprived (-Q) medium for 24 h. Data is presented as mean +/- SEM relative to WT, +Q mRNA levels for n = 3 replicates. N = 1 (*Egfr*), N = 2 (*Erbb2, Erbb3*) independent experiments. **E.** mRNA levels of soluble *Nrg1 Type I*, membrane-tethered *Nrg1 Type III,* and *Nrg2* in *Nf2*$^{-/-}$ SCs grown in complete (+Q), Nrg1-deprived (*Nrg2*) or glutamine-deprived (-Q) medium. Data is presented as mean +/- SEM relative to +Q mRNA levels for n = 3 replicates. **F.** Quantitation of basal ezrin intensity in *Nf2*$^{-/-}$ SCs stimulated with conditioned medium from glutamine-deprived (-Q) or Nrg1-deprived *Nf2*$^{-/-}$ SCs for 30 min.  Data points are shown for n = 12 (-Q) or n = 13 (+Q) cells. N = 2 independent experiments. All data are depicted as mean +/- SEM. All P values were calculated with unpaired two-tailed Student's t-test. **p<0.01, ***p<0.001.

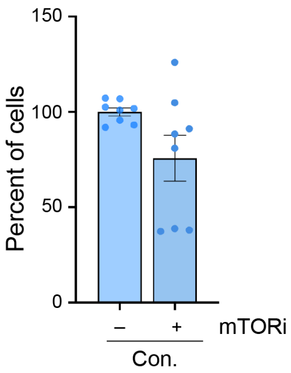


**Supplementary Figure 6.** Quantitation of cell viability of *Nf2*$^{-/-}$ SCs grown in complete medium upon treatment with mTORi (100 nM, 72h). Bars represent mean +/- SEM percentage of cells compared to vehicle treatment. n = 3, N = 2 independent experiments.

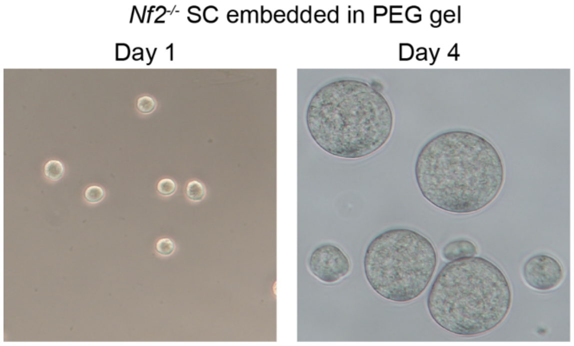


**Supplementary Figure 7.** Phase contrast images showing *Nf2*$^{-/-}$ SCs as single cells (Day 1) or spheres (Day 4) embedded in PEG gels.

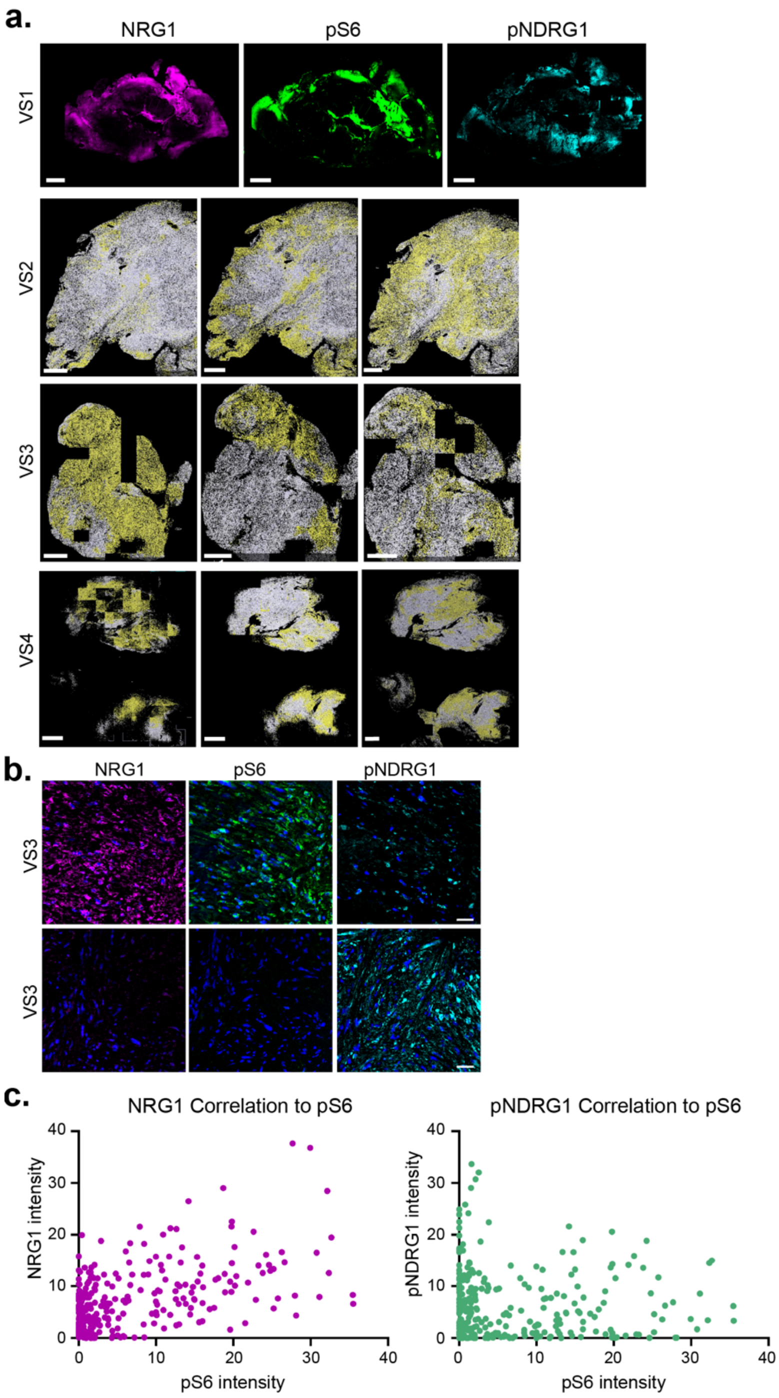


**Supplementary Figure 8. A.** *Top*, Immunofluorescence images showing NRG1 (magenta), pS6 (green) and pNDRG1 (cyan) staining across an entire tissue section from human vestibular schwannoma 1 (VS1). Lower, Segmentation analysis showing NRG1+, pS6+ or pNDRG1+ cells (all yellow) across entire sections of VS2, VS3 and VS4. Scale bars = 1 mm. **B.** Confocal images showing NRG1 (magenta), pS6 (green) and pNDRG1 (cyan) staining of aligned fields of view from VS3. Scale bar = 50 µm. **C.** xy scatter plots showing the positive correlation between Nrg1 and pS6 intensity and negative correlation between pNDRG1 and pS6 across fields of view from VS1, VS2, and VS3, related to Fig. 7 C,D (n = 16 ROIs in 5 20x fields of view per tumor).

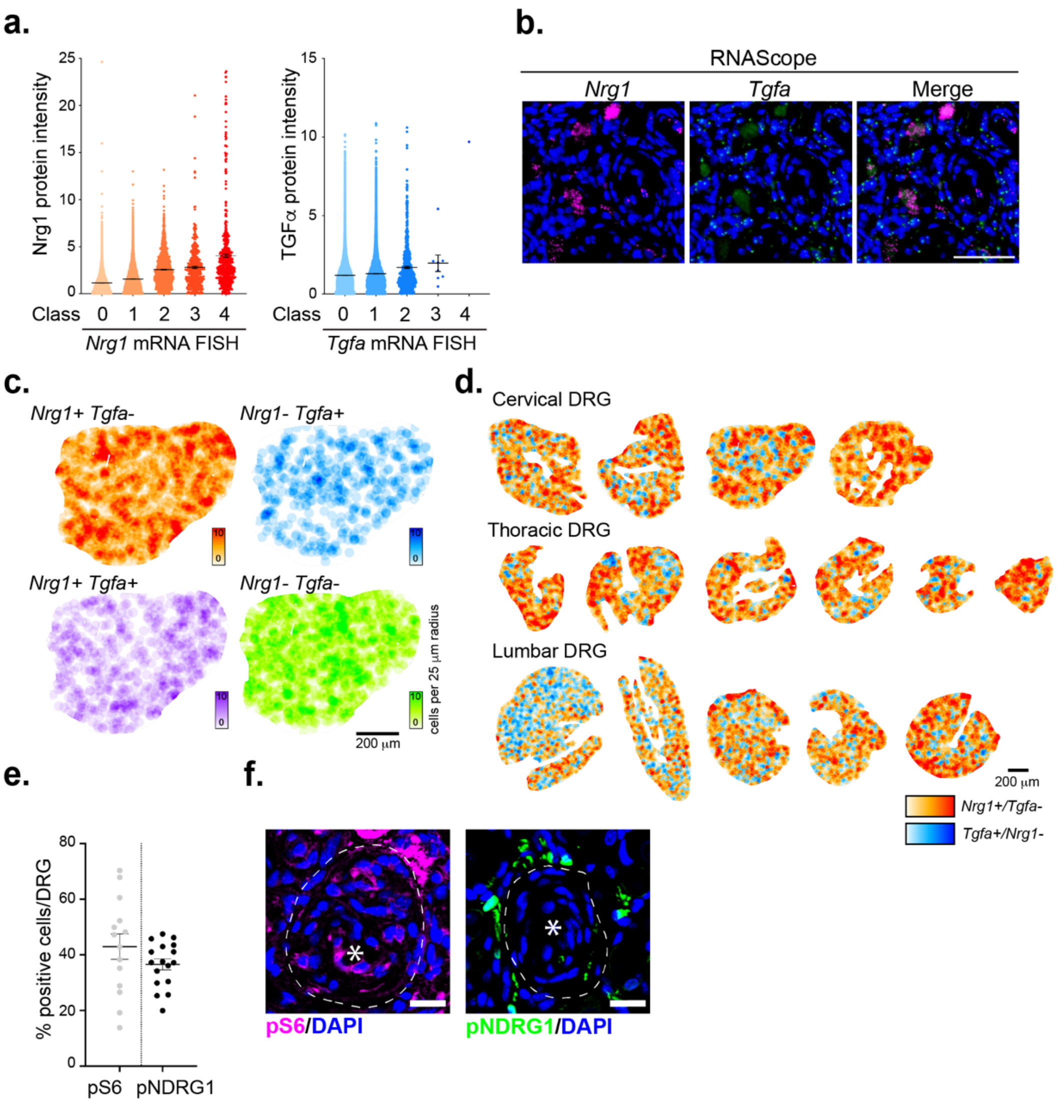


**Supplementary Figure 9. A.** Graphs showing the correlation between Nrg1 protein and *Nrg1* mRNA measured concomitantly by RNAscope. Data points show mRNA dots within each class as follows: Class 0 (0 dots/cell), Class 1 (1-3 dots/cell), Class 2 (4-9 dots/cell), Class 3 (10-15 dots/cell) or Class 4 (>15 dots/cell). **B.** Confocal images showing *Nrg1* (magenta), *Tgfa* (green) and merged mRNAs detected by FISH. Scale bar = 100 µm. **C.** Pseudocolored analysis of the distribution of *Nrg1+/Tgfa-*, *Nrg1/Tgfa+, Nrg1+/Tgfa+* and *Nrg1-/Tgfa-* mRNA expressing cells measured by FISH. Scale bar = 200 µm. **D.** A panel of DRG double pseudo-labelled for *Nrg1+/ Tgfa-* (orange) and *Tgfa+/Nrg1-* (blue) mRNA expressing cell populations. Scale bar = 200 µm.

**E.** Quantitation of pS6+ and pNDRG1+ cells across a panel of DRG from a *Postn-Cre;Nf2*$^{lox/lox}$ mouse. Each data point represents one DRG (n = 15, pS6; n = 18, pNDRG1). **F.** Confocal images showing the differential distribution of pS6+ cells within the whorls and pNDRG+ cells between whorls. Asterisks mark center of whorls. Scale bar = 20 μm. All data are depicted as mean +/- SEM.
